# A bacterial microbiome is dispensable for the induction of CD8 T cell exhaustion

**DOI:** 10.1101/2022.10.03.510696

**Authors:** Miriam Kuhlmann, Daphne Del Carmen Kolland, Gustavo Pereira de Almeida, Christian Hoffmann, Madlaina von Hoesslin, Jacqueline Berner, Christine Wurmser, Caspar Ohnmacht, Dietmar Zehn

## Abstract

Prolonged antigen exposure in chronic viral infections reduces the effector capacity of cytotoxic T cells - a phenomenon known as T cell exhaustion. Development of T cell exhaustion is driven by high viral titers, strong TCR stimulation, and high antigen concentrations associated with strong inflammatory signals. A largely unexplored factor has been the influence of the microbiome in these processes. Here, we report that T cell exhaustion progresses independently of the presence or absence of a microbiome in chronic lymphocytic choriomeningitis virus (LCMV) infections. Virus-specific CD8 T cells in germ-free mice showed high expression of the inhibitory receptor PD-1 and decreased cytokine production. Moreover, their global gene expression patterns, as determined by single-cell sequencing, were similar to those of cells in specific pathogen-free mice. In line with this, we observed similar pathogen loads with and without a microbiome. Thus, our study demonstrates that the microbiome is dispensable for the induction of T cell exhaustion and for the limited virus control seen in chronic LCMV infections.

## INTRODUCTION

Sustained antigen exposure, as occurs in chronic infections and tumors, triggers a differentiation program in T cells that is largely different from the program acquired by T cells during acute infection^1, 2^. Acute infections induce the proliferation and generation of large numbers of effector T cells that express high levels of effector molecules such as granzyme B, proinflammatory cytokines including tumor necrosis factor (TNF) and interferon gamma (IFNγ) while inhibitory receptors such as PD-1 and LAG-3 are only temporarily expressed. After infection has subsided, a population of memory cells can be detected in numbers largely exceeding the number of naïve T cells that were initially activated^3^. This memory population is maintained stably over long periods of time, and these cells mediate a strong recall response upon re-encounter with a similar or related pathogen. In contrast, although chronic infections also trigger strong activation and proliferation of pathogen-specific naïve T cells, the responding cells exhibit distinct phenotypic features including i) reduced expression of effector molecules like IFNγ and TNF, ii) stable expression of inhibitory receptors such as PD- 1, and iii) characteristic global gene expression profiles and epigenetic marks^2, 4^. Cells with this phenotype are typically referred to as exhausted T cells. These exhausted T cell populations exhibit cellular dynamics that are largely different from T cells in acute infections and chronic infections are ultimately not associated with the formation of a typical memory compartment.

Exhausted T cell populations constitute the type of T cells that are available to be recruited for therapeutic purposes against malignant tumors or chronic infection. While their effector capacity is restricted compared to the cells found in acute infection, it is well established that they still possess significant anti-tumor activity following their induced reactivation. Such reactivation is typically induced by blocking the inhibitory receptor-ligand systems that are expressed by exhausted T cells^5^. In fact, blocking PD-1/PD-L1 with targeting antibodies was an important proof of principle demonstrating that anti-tumor potential is contained in exhausted T cells. Moreover, this finding paved the road for the line of therapy that is now known as checkpoint blockade^6^.

Checkpoint blockade has proven to be extremely successful in many tumors compared to classical chemotherapy. Nevertheless, there are also a large number of patients in whom checkpoint blockade has limited or no effect. These failures suggest that improving activation of the immune system and overcoming the phenomenon of T cell exhaustion by other means than with currently available reagents hold substantial promise for more efficient therapies. A prerequisite for such developments is a deeper understanding of the molecular mechanisms and host factors that determine T cell exhaustion. A largely unexplored question has been how the bacterial microbiome affects the development and the degree of T cell exhaustion that occurs in chronic infections and our ability to overcome this phenotype. Indeed, numerous reports indicate that the microbiome is a major factor influencing adaptive immunity but its impact on the prototypic model system of T cell exhaustion - lymphocytic choriomeningitis virus (LCMV) infections - remained elusive^7^. Interestingly, we and others have noticed that chronic LCMV infection leads to massive yet transient changes in the bacterial microbiome during the course of infection.^8^ We therefore saw a need to systematically investigate, in well-controlled experimental systems, how the presence, or absence, or specific alterations of the bacterial microbiome affect the T cell response in chronic infections. Unexpectedly, we found that CD8 T cell exhaustion and pathogen control was barely affected by the absence of microbial colonization. These findings indicate that the microbiome does not affect phenotypic differentiation and effector function of virus-specific CD8 T cells in chronic infection.

## RESULTS

### Chronic infections induce massive but reversible changes in the bacterial microbiome

The intestinal microbiome shows a highly dynamic composition and is to a significant level constantly shaped and reshaped following bacterial exposure, alimentary alterations, antibiosis, or infections. We began our studies by looking at how the microbiome changes following exposure to acute and chronic LCMV infection - a prototypic model for studying T cell mediated immune responses in mice. The LCMV model has the advantage that several closely related strains exist that cause either acutely resolved infection or persisting chronic infection^7^. We used the LCMV armstrong strain, which causes an acute infection that is usually resolved within 5-10 days, and LCMV clone-13 or LCMV docile infections that persist for the life of the mice when applied at higher doses^9, 10^.

To determine how the microbiome is affected during these infections, we horizontally collected fecal samples from mice before and at different time-points after the infection with acute LCMV armstrong or chronic LCMV clone-13 (**Fig. 1A**). We noted only minor changes in the alpha-diversity shown by both the Richness and the Shannon effective following acute infection (**Fig. 1B**). In contrast, there were massive changes in chronic infection (**Fig. 1B**). In fact, the microbiome of mice chronically infected with LCMV displayed a reduction in alpha-diversity and a shift in beta-diversity 8 days post infection which was not observed in acute infection (**Fig. 1C**). Importantly, this shift was only temporary though and the microbiome reverted to its original state over time in chronic infection (**Fig. 1C**).

**Figure 1.**
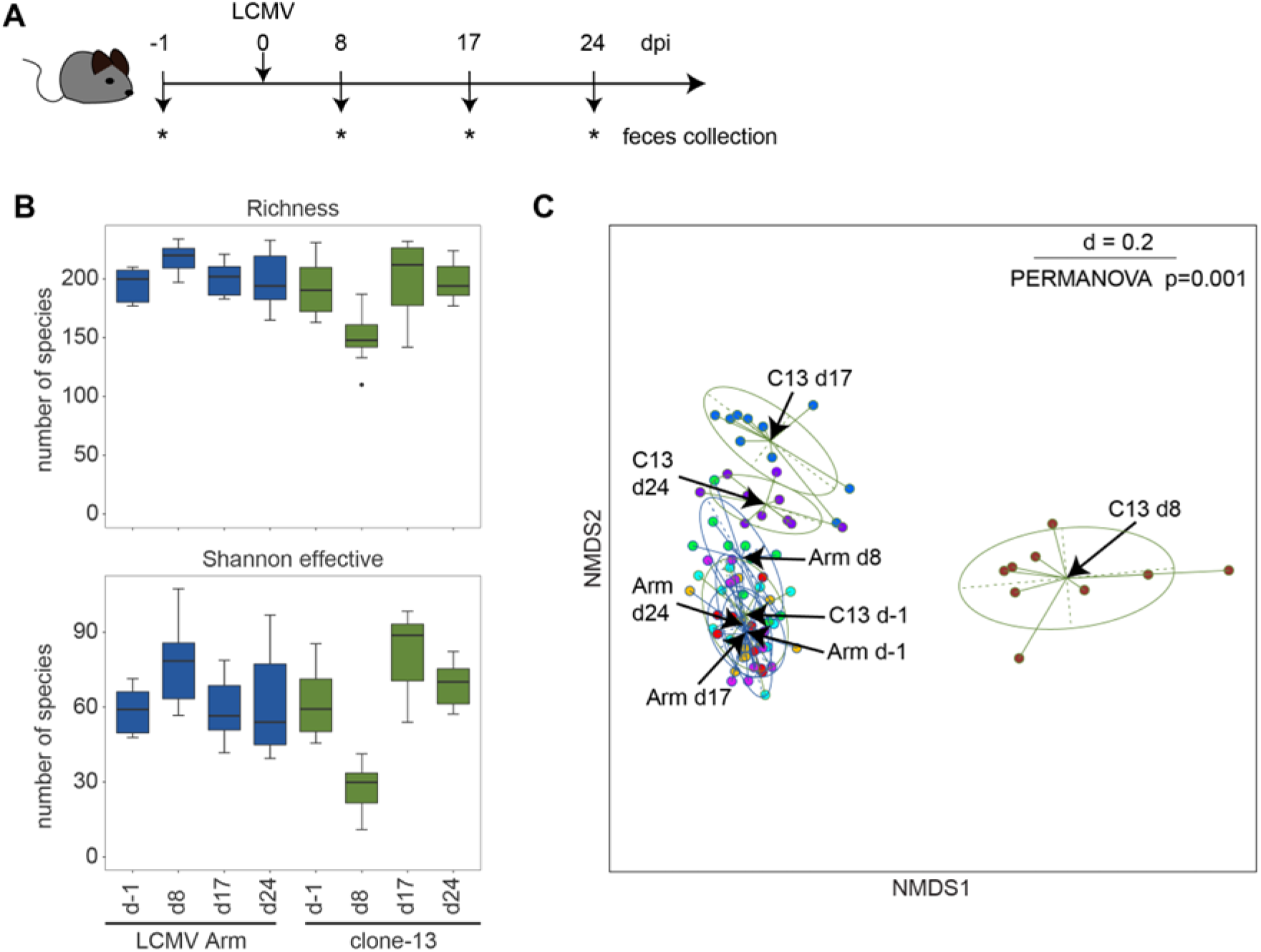
Microbiome dynamics during acute (armstrong) and chronic LCMV (clone-13) infections. SPF C57BL/6 mice were infected with LCMV armstrong (blue) or LCMV clone-13 (green). Stool samples were collected one day before and 8, 17, and 24 days after infection (dpi) as illustrated in (**A**). **B**) Plots represent the alpha-diversity of the microbiome of the different time-points as the richness and the Shannon effective number of species. **C**) The metaNMDS plot shows the beta-diversity of all microbiota in both infection models; dots represent individual mice. Samples are grouped according to the time-point at which they were taken.

### Similar expansion of virus-specific CD8 T cells in the spleen and small intestine in the absence of the microbiome

The strong changes in the microbiome over the course of the LCMV clone-13 infection raised our interest in determining if these changes impact the T cell response and in particular the T cell phenotype that arises during chronic infection. To assess the principal impact of the microbiome, we compared the response of T cells in specific pathogen-free (SPF) mice with mice that do not contain a bacterial microbiome (germ-free). We infected germ-free and SPF mice with LCMV clone-13 and used peptide-MHC tetramers loaded with the LCMV derived gp33 peptide to assess the response of virus-specific CD8 T cells in chronic LCMV clone-13 infections. We found that SPF and germ-free mice contained similar frequencies (**Fig. 2A**) but slightly reduced total numbers of gp33 tetramer-positive T cells in both the spleen and in the lamina propria of the small intestine (SI-LP) in germ-free mice at 8 and 28 days post infection (**Fig. 2B**). Of note, SPF mice also contained higher total CD8 T cells numbers (**Fig. 2C**). Hence, it is likely that the number of gp33-specific T cells in SPF mice are a consequence of higher total CD8 T cell numbers while the per cell expansion capacity appears similar. Thus, we did not find evidence for altered expansion with and without a bacterial microbiome.

**Figure 2:**
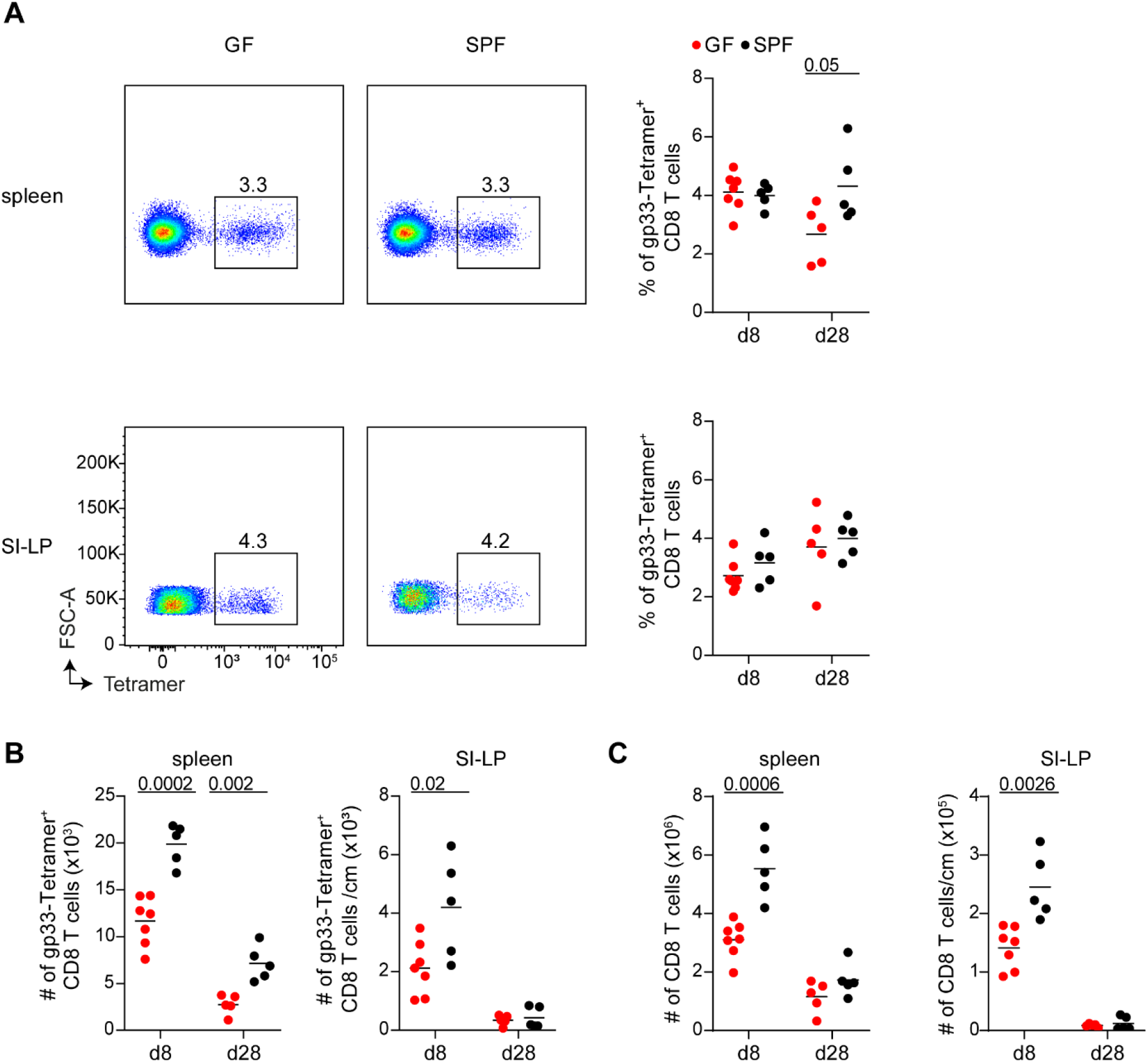
Similar expansion and maintenance kinetic of virus-specific CD8 T cells in chronic infections with and without the microbiome. Experimental design: Germ-free (GF) and SPF C57BL/6 mice were infected with LCMV clone-13 and analyzed on day 8 or day 28 after infection. **A**) Representative, CD8 gated flow cytometry plots showing the frequency of tetramer^+^ cells of germ-free and SPF mice on day 8 and d28 after infection in the spleen and the lamina propria of the small intestine (SI-LP). The dot plots on the right show the frequencies for all germ-free (red) and SPF (black) mice. **B**) Enumeration of total tetramer^+^ cells of germ-free and SPF mice on day 8 and d28 after infection in the spleen and per centimeter of the SI-LP. **C**) Enumeration of total CD8 T cells of germ-free and SPF mice on day 8 and d28 after infection in the spleen and per centimeter of the SI-LP. Colors as indicated in (A). All data are representative for at least two independently performed experiments with at least n ≥ 5, symbols indicate individual mice; horizontal lines show the mean. P values are from unpaired t-test.

Interestingly, we did not detect major changes in the phenotype of gp33-specific T cells with and without a microbiome. Co-staining of gp33 tetramer-positive T cells obtained from the spleen and the small intestine on day 8 and 28 post LCMV clone-13 infection revealed similar mean fluorescence intensities of PD-1 expression under all conditions (**Fig. 3A, B**). Similarly, cells from either SPF or GF-mice showed a comparable effector phenotype, as they lacked CD127 expression (**Fig. 3C**) and remained negative for KLRG1 (**Fig. 3D**). Of note, the absence of KLRG1 is a typical sign for exhausted effector T cells in chronic LCMV infection compared to cells found in acute and resolved LCMV infections, where a prominent KLRG1 expressing population can be detected^11^. Next, we also assessed the cytokine secretion profile of CD8 T cells. Following brief ex-vivo restimulation with gp33 peptide and subsequent intracellular staining, we detected similar IFNγ and TNF profiles in all conditions (**Fig. 4 and Supp. Fig. 1**). Consistent with established expression patters of exhausted T cells^11^, we noted that of the cytokine secreting cells, only a small fraction co-produced IFNγ and TNF while the majority of cytokine-positive cells produced only IFNγ and not TNF. Moreover, the IFNγ production of these cells was lower than of the IFNγ and TNF double producing cells (**Fig. 4**).

**Figure 3:**
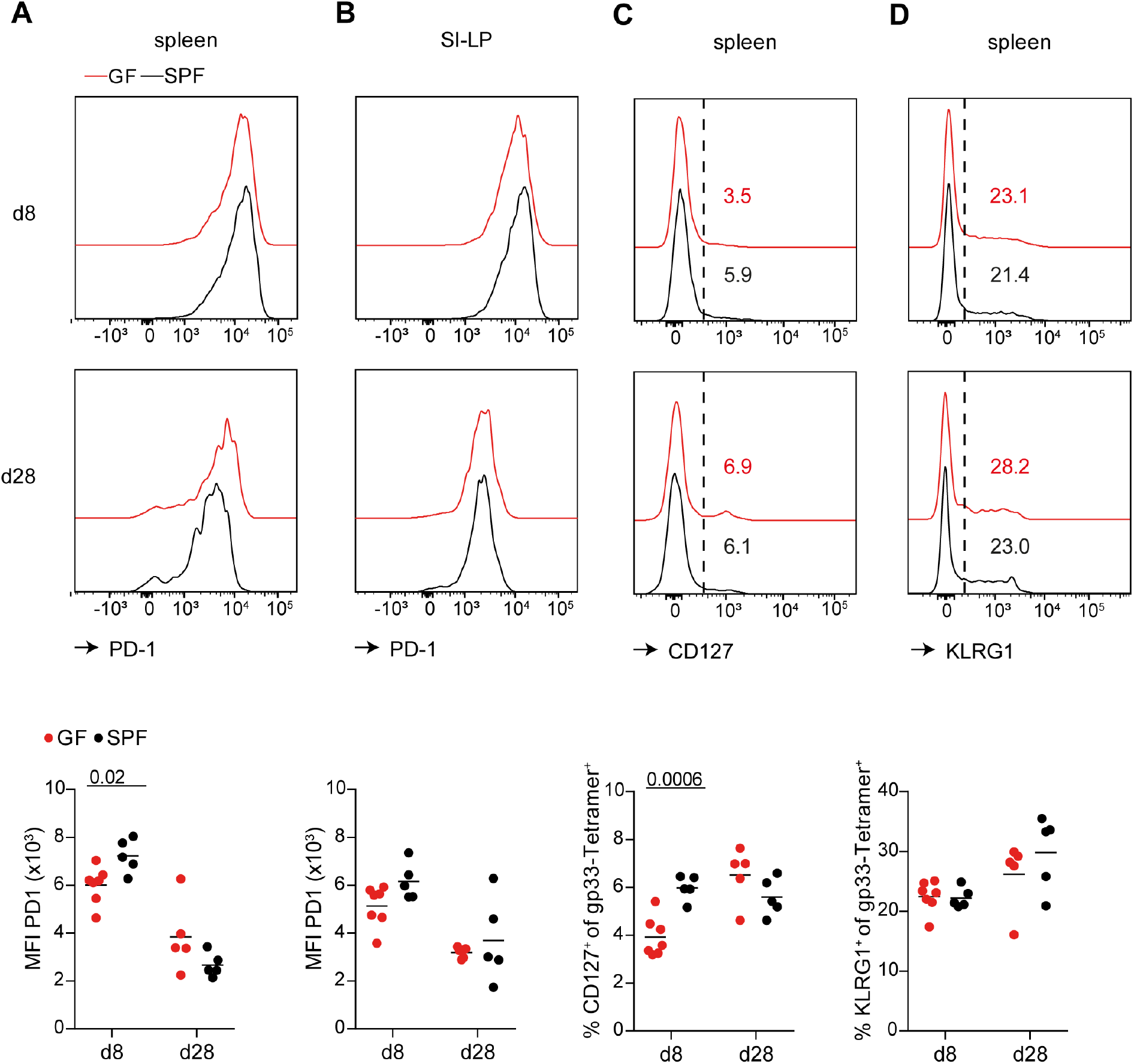
Similar phenotype of virus-specific CD8 T cells in chronic infections with and without the microbiome. Experimental design: Germ-free (GF) and SPF C57BL/6 mice were infected with LCMV clone-13 and analyzed on day 8 or day 28 after infection. **A** and **B**) Representative, tetramer^+^ gated histograms showing the expression of PD-1 for germ-free and SPF mice on day 8 and d28 after infection in the spleen (A) and SI-LP (B). The dot plots at the bottom show the MFI of PD-1 on tetramer^+^ cells for all germ-free and SPF mice. **C**) Representative, tetramer^+^ gated histograms showing expression of CD127 for germ-free (red) and SPF (black) mice on day 8 and d28 after infection in the spleen. The dot plot at the bottom shows the frequencies of CD127^+^ cells for all germ-free and SPF mice, Color coding as indicated in (A). **D**) Representative, tetramer^+^ gated histograms showing the expression of KLRG1 for germ-free and SPF mice on day 8 and d28 after infection in the spleen. The dot plot at the bottom shows the frequencies of KLRG1^+^ cells for all germ-free and SPF mice. Colors as indicated in (A). All data are representative for at least two independently performed experiments with at least n ≥ 5, symbols indicate individual mice; horizontal lines show the mean. P values are from unpaired t-test.

**Figure 4:**
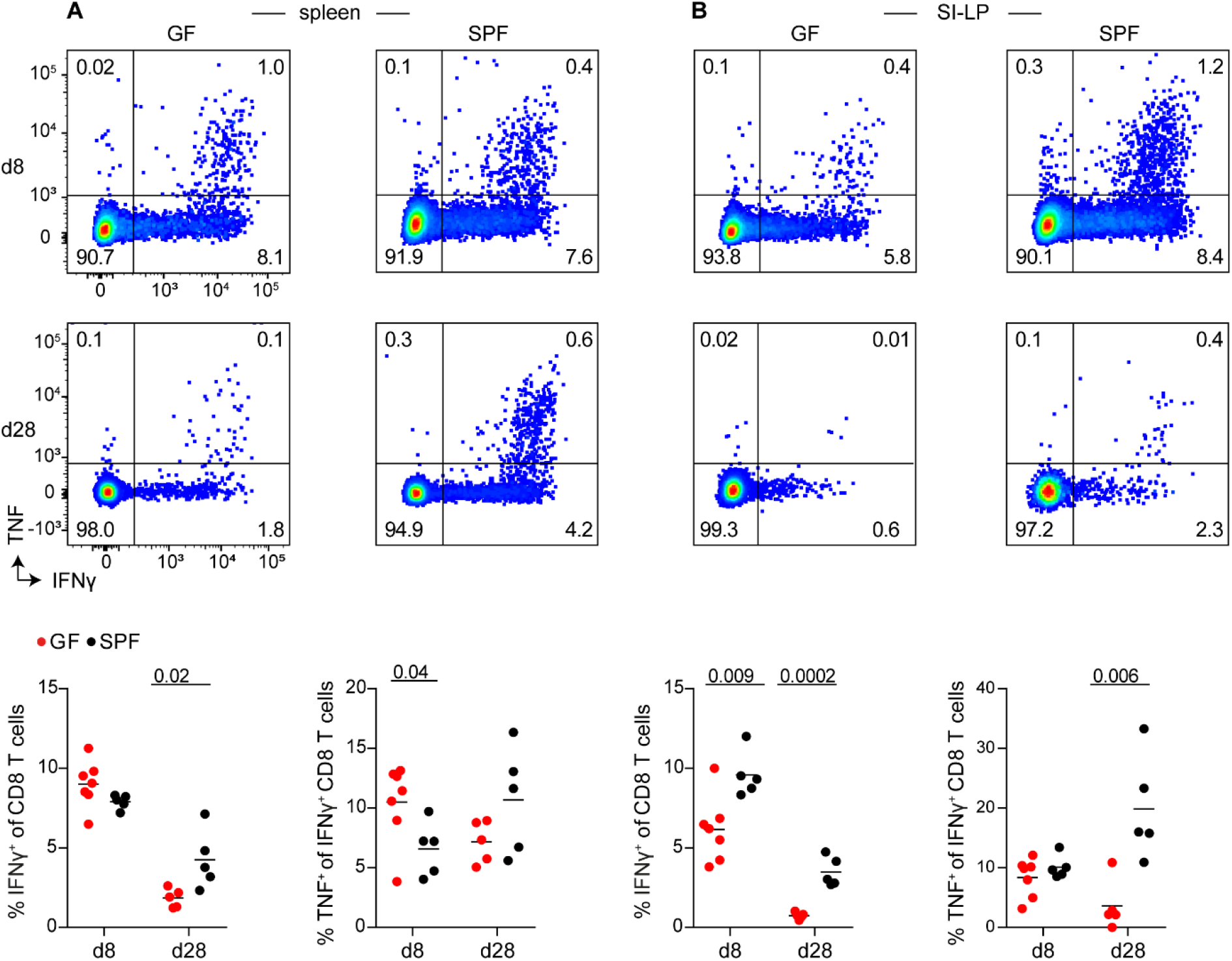
Absence of a microbiome reduces the cytokine secretion capacity of CD8 T cells in the intestine. Experimental design: Germ-free (GF) and SPF C57BL/6 mice were infected with LCMV clone-13 and analyzed on day 8 or day 28 after infection. **A** and **B**) Representative, CD8 gated flow cytometry plots showing the frequency of IFNγ and TNF expressing cells of germ- free and SPF mice on day 8 and d28 after infection after restimulation with gp33-41 in the spleen (A) and the SI-LP (B). The dot plots at the bottom show the frequencies of IFNγ expressing (left) CD8 T cells and frequencies of TNF^+^ of IFNγ expressing (right) CD8 T cells for all germ-free (red) and SPF (black) mice. All data are representative for at least two independently performed experiments with at least n ≥ 5, symbols indicate individual mice; horizontal lines show the mean. P values are from unpaired t-test.

Both strains, LCMV clone-13 and LCMV docile, induce chronic infections but the level of virus persistence and the level of T cell exhaustion is higher in LCMV docile compared to clone-13 infections. We therefore repeated our experiments using chronic LCMV docile infections. We noted a small tendency towards a slightly higher frequency of IFNγ producing cells in the lamina propria of SPF mice but, overall, the phenotype of the T cells was again very similar in SPF and germ-free mice (**Fig. 5**). **We therefore conclude that LCMV-specific T cells acquire similar phenotypic signs of T cell exhaustion irrespectively of the presence of a bacterial microbiome.**

**Figure 5:**
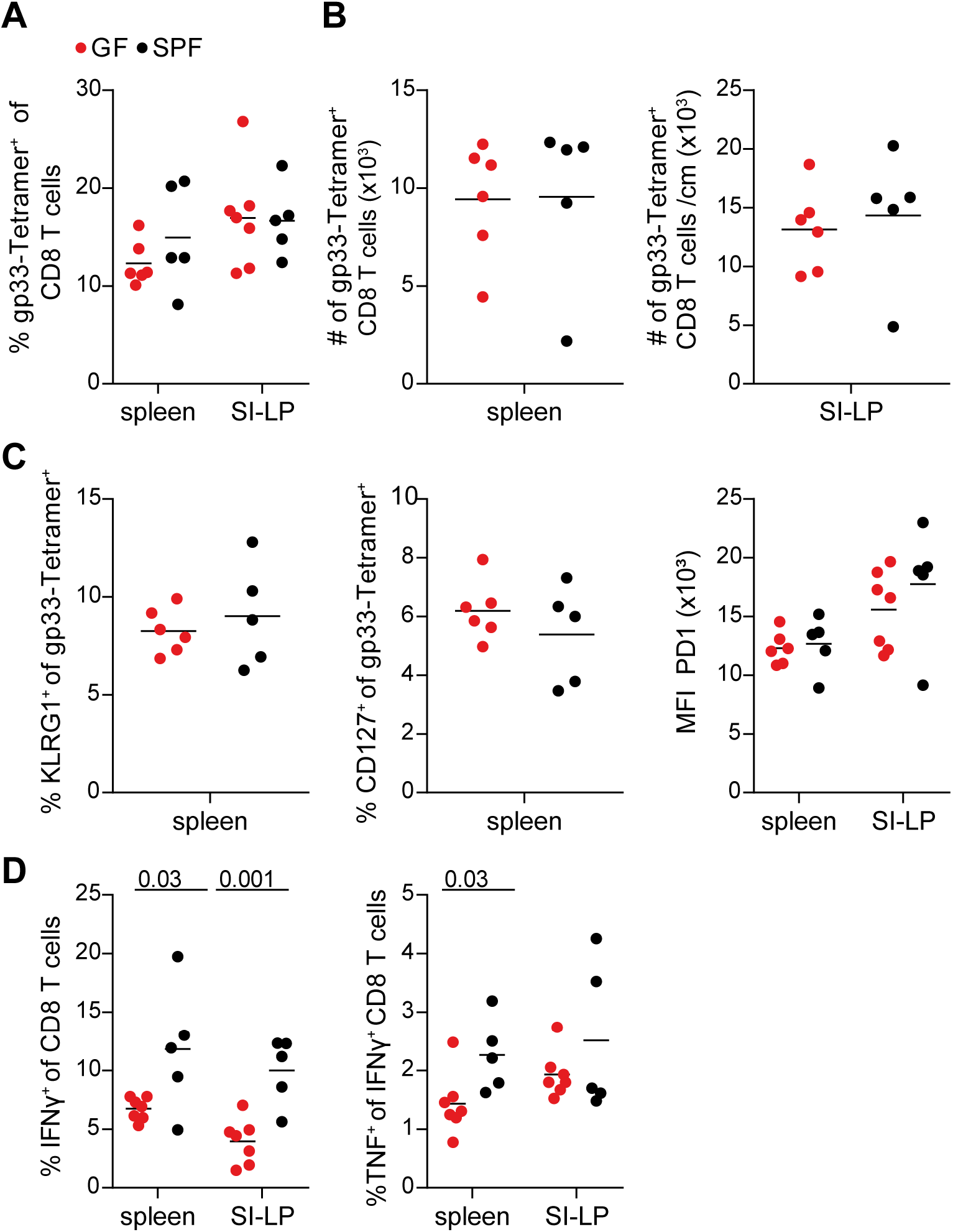
Kinetic and phenotype of virus-specific CD8 T cells is independent of the microbiome in LCMV docile infections. Experimental design: Germ-free (GF) and SPF C57BL/6 mice were infected with LCMV docile and analyzed on day 16 after infection. **A**) Frequencies of tetramer^+^ cells of germ-free (red) and SPF mice (black) in the spleen and SI-LP. **B**) Numbers of tetramer^+^ cells of germ-free and SPF mice in the spleen (left) and per centimeter of the SI- LP (right). Colors as indicated in (A). **C**) Frequencies of KLRG1^+^ and CD127^+^ and MFI of PD-1 of tetramer^+^ cells for germ-free and SPF mice in the spleen (KLRG1, CD127 and PD-1) and the SI- LP (PD-1). Colors as indicated in (A). **D**) Frequencies of IFNγ expressing (left) CD8 T cells and frequencies of TNF^+^ of IFNγ expressing (right) CD8 T cells for germ-free and SPF mice. Colors as indicated in (A). All data are representative for at least two independently performed experiments with at least n ≥ 5, symbols indicate individual mice; horizontal lines show the mean. P values are from unpaired t-test.

### LCMV-specific CD8 T cells show similar global gene expression profiles in germ-free and SPF mice

To explore more broadly the impact of the microbiome on LCMV specific CD8 T cells, we decided to characterize these cells more thoroughly using an unbiased approach. We therefore collected ∼1000-5000 gp33 tetramer-positive cells sorted from the lamina propria of the small intestine and from the spleen of three germ-free and three SPF mice and performed single cell-resolved RNA sequencing (**Fig. 6A**). A merged 2-dimensional UMAP representation of all samples is shown in **Figure 6B**. By unsupervised KNN clustering, we found two clusters specific for the spleen, three for the small intestine, while three clusters contained cells from both organs (**Fig. 6C**). Based on the expression of marker genes which are uniquely up- or downregulated in a specific cluster, we made the following assignments: Cluster S1 = circulating effector T cell cluster (i.e., *Klf2*⇑, *Cxcr3*⇑, *S1pr1*⇑), Cluster S2 = progenitor T cell cluster (i.e., *Tcf7*(TCF1)⇑, *Id3*⇑, *Sell*⇑, *Cxcr5*⇑, *Ccr7*⇑, *GzmB*⇓ not shown, ns), Cluster I1 = intestinal effector T cell cluster with signs of T cell exhaustion (*Itgae*[CD103])⇓, *Cxcr6*⇑ns, *Pdcd1*⇑ns, *Lag-3*⇑ns), Cluster I2 = intestinal effector T cell cluster (*Itgae*[CD103]⇓, *Cd69*⇑ns, *Xcl1*⇑, *Icos*⇑ns, *Batf*⇑ns, *Nr4a1*⇑, *Nr4a2*⇑ns, *Nr4a3*⇑), Cluster I3 = tissue resident intestinal T cells (*Itgae*[CD103]⇑), Cluster C1 = possibly dying cells (rich in mitochondrial genes), Cluster C2 = proliferating cluster (i.e., *Mki67*⇑, cell cycle genes⇑), Cluster C3 = effector T cell cluster (multiple effector T cell genes⇑) (**Fig. 6D**). T cells from SPF and germ-free mice had a similar percentage distribution among these clusters (**Fig. 6E**). **We conclude therefore that T cells are able to acquire an exhausted phenotype even in germ-free mice. This corroborates our conclusion that the microbiome has a limited impact on the phenotype of CD8 T cells during LCMV infections.**

**Figure 6:**
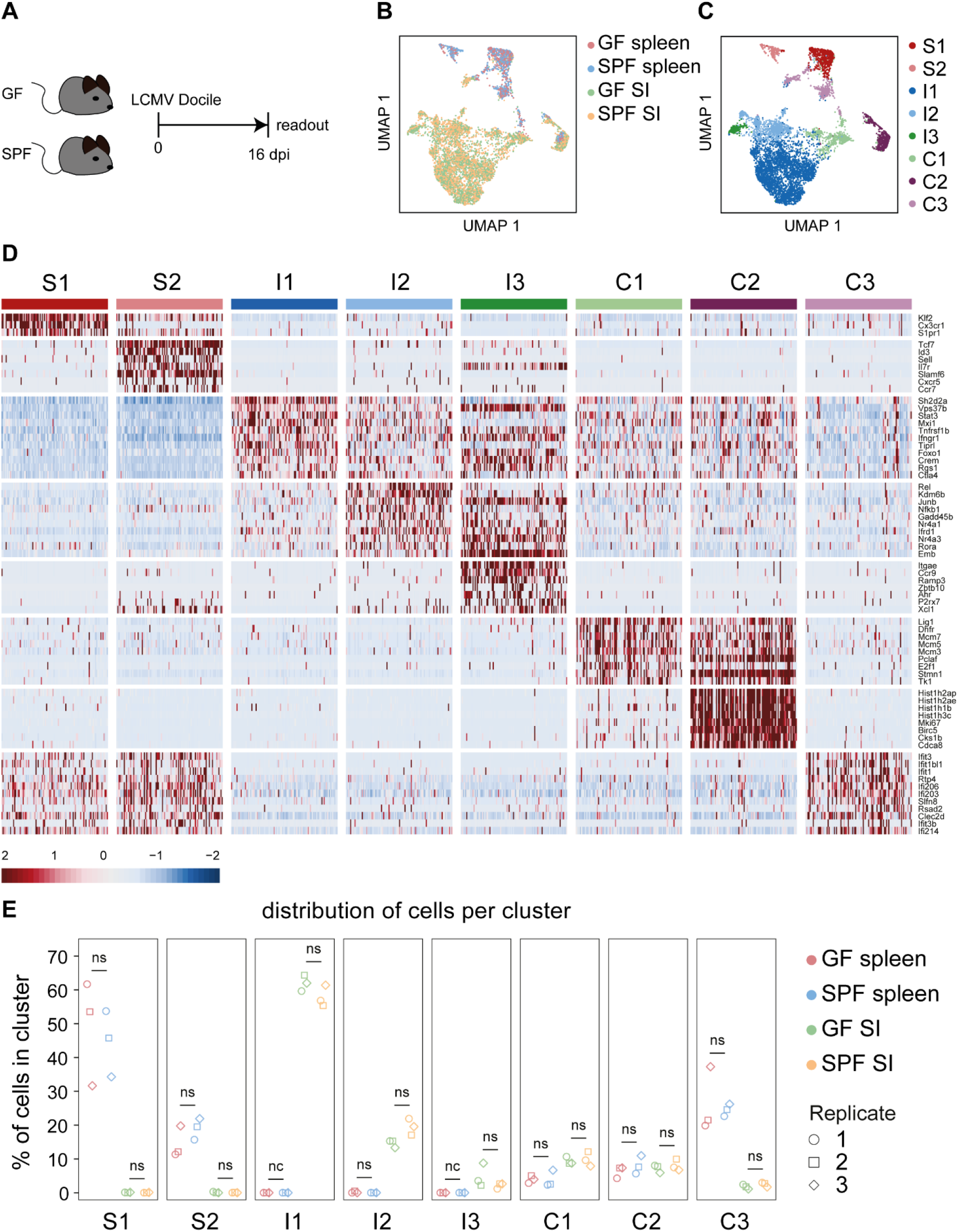
Single-cell RNA sequencing reveals a limited impact of the microbiome on the transcriptional profile of virus-specific CD8 T cells. **A**) Experimental design: Germ-free (GF) and SPF C57BL/6 mice were infected with LCMV docile and tetramer^+^ CD8 T cells were sorted 16 days post infection from the spleen and SI-LP. **B**) Uniform Manifold Approximation and Projection (UMAP) showing the distribution of cells originating from the spleen (SP) and small intestine lamina propria (SI) in GF and SPF mice. **C**) UMAP showing the clusters identified by unsupervised KNN: 2 clusters specific for the spleen (S), 3 clusters specific for the small intestine (I), and 3 common clusters (C). **D**) Heat map illustrates z-scores of genes of interest selected from the top-20 marker genes of the individual clusters. **E**) Plots show the percentage wise distribution of cells of all analyzed samples into these clusters.

### Similar pathogen control after chronic LCMV infections in the absence of the microbiome

Next, we tested the pathogen control and effector function of CD8 T cells in germ-free and SPF mice. We noted no impairment of virus control in germ-free compared to SPF mice. The level of plaque-forming units (PFU) was comparable in the spleen, liver, kidney, and small intestine 16 days post LCMV docile infection (**Fig. 7**).

**Figure 7:**
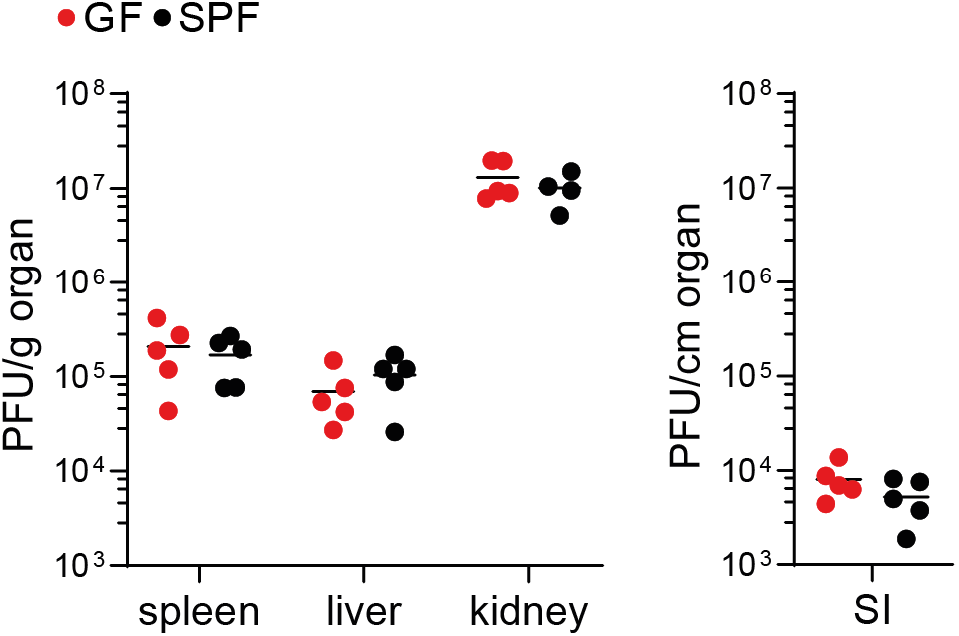
Absence of the microbiome significantly reduces virus control in chronic LCMV infections. Experimental design: Germ-free (GF, red) and SPF (black) C57BL/6 mice were infected with LCMV docile. Viral titers in the spleen, liver, kidney (PFU/g), and SI (PFU/cm) were assessed on day 16 after infection. All data are representative for at least two independently performed experiments with at least n ≥ 4, symbols indicate individual mice; horizontal lines show the mean.

The complete absence of microbes and all their metabolites and cues in germ-free mice represents a unique situation associated with several anomalies particularly in the immune system^12^. To explore the impact of the microbiome in a less extreme setting, we repeated our infection experiments to include a group of mice that contained only the minimum oligo-mouse-microbiota (oligoMM-12). This consortium includes 12 strains representing members of the major bacterial phyla in the murine gut. These are *Acutalibacter muris* KB18, *Flavonifractor plautii* YL31, *Enterocloster clostridioforme* YL32, *Blautia coccoides* YL58, *Clostridium innocuum* I46, *Limosilactobacillus reuteri* I49, *Enterococcus faecalis* KB1, *Bacteroides caecimuris* I48, *Muribaculum intestinale* YL27, *Bifidobacterium longum subsp. animalis* YL2, *Turicimonas muris* YL45 and *Akkermansia muciniphila* YL44. The consortium was developed initially as a stable minimal consortium^13^. Upon LCMV clone-13 infection, we confirmed that CD8 T cells have a similar phenotype in the different types of hosts including the oligoMM-12 mice (**Supp. Fig. 2A-D**), again corroborating the idea that CD8 T cell exhaustion develops independent of the microbiome.

We noted a minor impairment of virus control in spleen, liver, and kidney in both germ-free and oligoMM-12 mice compared to SPF mice (**Supp. Fig. 2E**) on day 28 pi. GF mice contained in the small intestine titer of about 10^3^ Pfu per cm organ, while SPF mice were free. In the oligoMM-12 group, 3 out of 5 mice were negative while two mice showed titer similar to the germ-free mice. Titers of ∼10^3^ PFU are very low and close to the limit of detection – which likely explained the mixed response in the oligoMM-12. **Overall, we conclude therefore that the bacterial microbiome is dispensable for the generation and modulation of an exhausted phenotype and does only have a transient and local impact on the level of pathogen control in chronic LCMV infection.**

## DISCUSSION

We have addressed how the absence of a bacterial microbiome impacts the formation and the phenotype of exhausted T cells. Unexpectedly, using flow cytometry and deep transcriptional expression profiling, we observed that T cells primed by chronic LCMV docile or LCMV clone-13 infections presented themselves with an identical phenotype and similar relative subset distribution regardless of the presence of a microbiome. Interestingly, this outcome contrasts with observations in which alterations in the composition of the microbiome went along with a strong impact on T cell phenotypes and effector function mediated by them. For instance, it was shown that a specific consortium of bacterial commensals can elicit stronger IFNγ induction in the CD8 T cell compartment, which correlated with enhanced host resistance to *Listeria monocytogenes* infection and improved checkpoint inhibition efficacy in a tumor model^14^. Similarly, commensal bacteria were shown to stipulate first line defense against Listeria infections^15^. Moreover, the different types of infections in so called “dirty mice” showed enhanced resistance to microbial challenge further underlining that alterations in the microbiome and unrelated prior pathogen-challenges increase protective immunity^16, 17^. While these studies underscored the potential of the microbiome to profoundly impact CD8 mediated protective immunity and to alter vaccine responses, key aspects of the mechanisms underlaying these beneficial effects remain poorly understood. Here, we observed however that the microbiome has no effect on the CD8 T cell response during chronic LCMV infection. This suggests that the inflammatory environment present during LCMV infection dominates over preexisting and concomitant microbial-derived signals present or absent in SPF or germ-free mice, respectively. LCMV infections are associated with high production of type 1 interferons and other proinflammatory cytokines, which likely create an environment that determines T cell phenotype and function. In line with this, enteric infections with murine norovirus (MNV) in germ-free mice have been shown to Iead to a normalization of intestinal morphology and local immune function. Strikingly, the ability of MNV to compensate for commensal bacteria in this setting were critically depended on IFN I signalling^18^. Of note, our mice are free from norovirus infection.

There are several other studies that have explored links between LCMV infections and the microbiome but these focused predominately on the effect of the infection on the microbiome^8, 19, 20^. Moreover, there is a historic study describing that intracerebral LCMV infection resulted in better survival of germ-free mice compared to conventional mice^21^. These data are not easy to interpret as the intracerebral infection is different from the routes that are typically used to study acute and chronic LCMV infection but also because this particular publication lacks insights into the T cell biology. However, it needs to be considered that LCMV is not a cytopathic virus such that high titers are tolerated by mice in the absence of T cells. In fact, immunopathology and lethal outcomes of LCMV infection are typically associated with too strong T cell responses^22^ and that these additional T cells might be sufficient to drive pathology while eliminating the virus.

We were also surprised about outcomes we observed in the absence of a microbiome if one considers prior reports that structure of secondary lymphoid organs are underdeveloped or differentially organized in the absence of a microbiome^23–25^. For unclear reasons these deficits did not impair the ability to prime a profound CD8 T cell response to LCMV and for inducing T cell exhaustion. Whether virus-specific CD4 T helper cell responses are affected by microbial colonization is part of ongoing investigations, but preliminary results suggest a more profound effect of the microbiome on this compartment. Moreover, microbiota-dependent regulatory T cells constitute a cellular subset that links the microbiome to the regulation of immune responses during infections^26^. Such peripherally induced Treg cells, generated during a critical time window around weaning along with other microbiome- responding cell types, have also been proposed to counteract the general immune tone in germ-free mice from a type 2 bias and associated risk of atopic disorders to a more equilibrated immune system^27–29^. During chronic LCMV infection, however, absence of microbiome-induced Treg cells does not seem to affect the virus-specific CD8 T cell response.

We have started this project after moving to a new animal facility in which the mice showed alteration in the exhausted T cell phenotype such as slightly lower PD-1 expression levels and mildly upregulated cytokine secretion. Since we were still using the same mouse colony including similar in-house bread C57BL/6 host mice and similar food, we reasoned that alterations in the microbiome could be the reason. In particular, findings that differences in animal husbandry lead to significant changes in the microbiome raised our interest in this topic^30^. Similarly, the microbiome after antibiotic treatments was also shown to massively alter the outcome of CD8 T cells during viral infection^31, 32^. We thus suspected that a distinct microbial composition in the different animal facilities might have led to the phenotypic differences we have observed. Based on these observations as well as the fact that viral control in germ-free mice is transiently impaired on a low level, one could speculate that microbially-derived signals could play at least a limited role in fine-tuning the extent to which exhaustion manifests itself as well as fine-tuning antiviral effector functions. Future experiments would be required to explore these possibilities in detail.

Overall, our data show however that the phenomenon of CD8 T cell exhaustion can be readily induced in germ-free animals during chronic LCMV infection irrespectively of the microbiome.

## MATERIAL AND METHODS

### Mice

Specific pathogen-free (SPF) C57BL/6 mice were purchased from Charles River, germ-free (GF) C57BL/6 mice were provided by the Central Animal Facility (Hannover Medical School, Germany) and the Core Facility Gnotobiology (ZIEL - Institute for Food & Health, Technical University of Munich, Germany), and oligoMM-12 C57BL/6 mice were obtained from the Central Animal Facility (Hannover Medical School, Germany). P14^33^ mice were provided by Annette Oxenius (ETH Zürich, Switzerland).

SPF mice were bred and maintained in specific pathogen-free facilities and infected in modified specific pathogen-free animal facilities at the Technical University of Munich (Germany). GF and oligoMM-12 mice were housed and infected in open cages within plastic film isolators and located in a room with a controlled environment at the Technical University of Munich (Germany).

Experiments were carried out in male or female mice that were at least six weeks old in strict accordance with the institutional and governmental regulations in Germany. All procedures were approved by the responsible veterinarian authorities of the Regierung von Oberbayern in Germany. Animals were randomly assigned to experimental groups which were non-blinded and no specific method was used to calculate sample sizes.

### Infections

LCMV stocks were propagated in BHK-21 cells (provided by M. J. Bevan) cultured at 37 °C in Dulbecco’s Modified Eagle Medium (DMEM) supplemented with 10% fetal calf serum (FCS). Frozen LCMV stocks were diluted in PBS (Gibco) and 2×10^5^ plaque-forming units (PFU) LCMV armstrong were injected intraperitoneally (i.p.), 5×10^6^ PFU LCMV clone-13 or 2×10^4^ PFU LCMV docile were injected intravenously (i.v.).

### Measurement of viral burden

To determine virus load, tissues were harvested, weighted, and suspensions were shock-frozen and stored at -80 °C to release the virus. Levels of virus were determined by plaque forming assay^11^ using Vero cells (provided by M. J. Bevan).

### Isolation of lymphocytes from tissues

Spleens were dispersed into single-cell suspensions by mashing through a 100-μm nylon cell strainer (BD Falcon). Red blood cells were lysed with a hypotonic ammonium-chloride-potassium buffer.

To obtain SI-LP lymphocytes, residual adipose tissue and Peyer’s patches were removed from the tissue. The small intestine was incubated in PBS containing 0.5 M EDTA for 30 min at 4 °C and afterwards washed in PBS and cut into small pieces. Digestion was performed by shaking in RPMI-medium (Gibco) with 240 mg/ml Liberase TL (Roche) and 150 μg/mL DNase I (Roche) at 37 °C for 60 min. Single-cell suspensions were obtained by filtering through a 100-μm nylon cell strainer. Cells were resuspended in 44% percoll (Cytiva) and underlaid with 67% percoll for density gradient centrifugation.

### Surface and intracellular antibody staining and flow cytometry cell sorting of mouse T cells

Blocking of unspecific antibody binding was achieved with 2.4G2 (BioXCell) and dead cells were stained using the Zombie NIR™ Fixable Viability Kit (Biolegend). Cells were stained with APC-conjugated MHC class I tetramers of H-2D^b^ complexed with LCMV gp33-41 (TCMetrix) for 1 h at 4 °C. Surface staining was performed for 30 min at 4 °C in PBS supplemented with 2% FCS (Sigma-Aldrich) and 0.01% azide (Sigma-Aldrich) (FACS-buffer) using the following antibodies. Anti-CD45.1 (A20), anti-KLRG1 (2F1), anti-CD127 (A7R34), and anti-PD-1 (J43) were purchased from eBioscience, and anti-CD8a (53-6.7) was obtained from Biolegend. Cells were fixed in PBS containing 2% formaldehyde for 15 min, then washed and resuspended in FACS-buffer. For intracellular cytokine staining, ∼3x10^6^ splenocytes were cultured in the absence or presence of gp33-41 for a total of 5 h at 37 °C. After 30 min, 7 μM brefeldin A was added. Following staining for surface antigens and fixation as described above, cells were permeabilized in PBS with 0.25% Saponin and 0.25% BSA (Perm-buffer). Staining with anti- IFNγ (XMG1.2) and anti-TNF (MP6-XT22) antibodies from eBioscience was performed for 40 min at 4 °C in Perm-buffer. For flow cytometry-based sorting, splenic and SI-LP cells were enriched for CD8 T cells using the mouse CD8α^+^ T cell isolation kit (Miltenyi Biotech). Virus-specific CD8 T cells were obtained by sorting of gp33-Tetramer^+^ CD8 T cells using a FACSAria Fusion instrument (BD). Flow cytometry measurements were conducted on an LSR-Fortessa (BD). Data were analyzed using FlowJo (TreeStar).

### Single-cell RNA sequencing and analysis

Gene expression libraries for single cell RNA sequencing were prepared by using the Chromium Next GEM Single Cell 3’ Reagent Kit v3.1 and the 10x Chromium Controller (10x Genomics) following the manufacturer’s protocol (CG000206 Rev D). Single Index Kit T Set A was used for multiplexing (i7 index read, 8bp). The samples were sequenced in a paired-end run (read 1: 28bp, read 2: 91bp) on a NovaSeq6000 platform using S1 v1.5 (100 cycles) sequencing kits (Illumina). Bcl2fastq software (v2.20.0.422) was used for demultiplexing and generation of .fastq files allowing zero barcode-mismatches. Alignment of sequencing reads and gene counting with single-cell resolution were performed with 10x Genomics Cell Ranger v6.0.1^34^, using the default parameters and the pre-built mouse reference v2020-A (10x Genomics, Inc) based on mm10 - GENCODE vM23/Ensembl 98. Only cells with more than 1200 genes detected, less than 10% of mitochondrial genes and with UMI counts less than 3 standard deviations above the mean were kept for downstream analysis.

Only data for genes detected in at least 3 cells in one of the samples were kept for further analysis. Contaminating cells were filtered out based on the average cluster expression of the marker genes Cd14, Lyz2, Fcgr3, Ms4a7, Fcer1g, Cst3, H2-Aa, Ly6d, Ms4a1, Ly6d, Cd19, Cd3e, Cd8a and mitochondrial genes. Raw read counts data from different tissues of the same donor were merged and normalized together using the R package sctransform v0.3.2^35^ (with glmGamPoi method). Downstream analysis was performed with the R package Seurat v4.0.1^36^. Anchors between replicates were identified on top 1000 highly variable genes and integration performed on first 20 dimensions. PCA was calculated on the top 1000 highly variable genes, KNN graph and UMAP were computed on the 30 nearest neighbors and first 20 PCA dimensions. Clusters were identified with the Louvain algorithm at resolution 0.36.

### DNA extraction and 16S rRNA gene sequencing

Fecal samples were collected on dry ice immediately after sampling and stored at -80 °C. DNA was extracted with a modified protocol by Godon et al.^37^ as previously described^38^. DNA was purified with the NucleoSpin gDNA Clean- up Kit (Machery-Nagel) following the manufacturer’s protocol and DNA concentration determined by using a NanoDrop. The 16S rRNA gene was amplified by PCR using barcoded primers flanking the V3 and V4 hypervariable regions (341F, 785R)^39, 40^. Sequencing was performed on the Illumina MiSeq platform according to the manufacturer’s instructions. Demultiplexed FASTQ files were transformed into zero-radius Operational Taxonomic Unit (zOTU) tables using the Integrated Microbial Next Generation Sequencing platform^41^ (trimming of ten nucleotides at the 5’ and 3’ end, abundance cutoff of 0.25%). Downstream analysis was performed using the R pipeline Rhea^42^. Alpha-Diversity was calculated as species richness and Shannon effective number of species. Distance matrix for beta-diversity was calculated based on the generalized UniFrac approach^43^. Statistical analysis was performed by permutational multivariate analysis of variance (PERMANOVA).

### General data analyses

Statistical analyses were performed with Prism 9.0 (Graphpad Software). Mann-Whitney test and unpaired t-test were used according to the type of experiment. P value < 0.05 was considered significant; p value > 0.05 was considered not significant.

## ACKNOWLEDGMENTS

We thank C. Amette, B. Dötterböck, and W. Schmid for technical assistance; C. Lechner for animal husbandry and the SFB1371 for providing the infrastructure to perform gnotobiotic experiments.

## AUTHOR CONTRIBUTION

Conceptualization: DZ, MK, CO; Formal analysis: MK, GPDA, MVH, DCK; Funding acquisition: DZ and CO; Investigation: MK, DCK, MVH, JB, CW; Methodology: MK; Visualization: MK, DZ; Writing: DZ, MK, CO

## DATA AND MATERIALS AVAILABILITY

All data will be made available upon publication. Next-generation sequencing data for this study have been deposited in the NCBI GEO repository with accession number XXXX.

