## Supplementary material for "A bacterial microbiome is dispensable for the induction of CD8 T cell exhaustion"

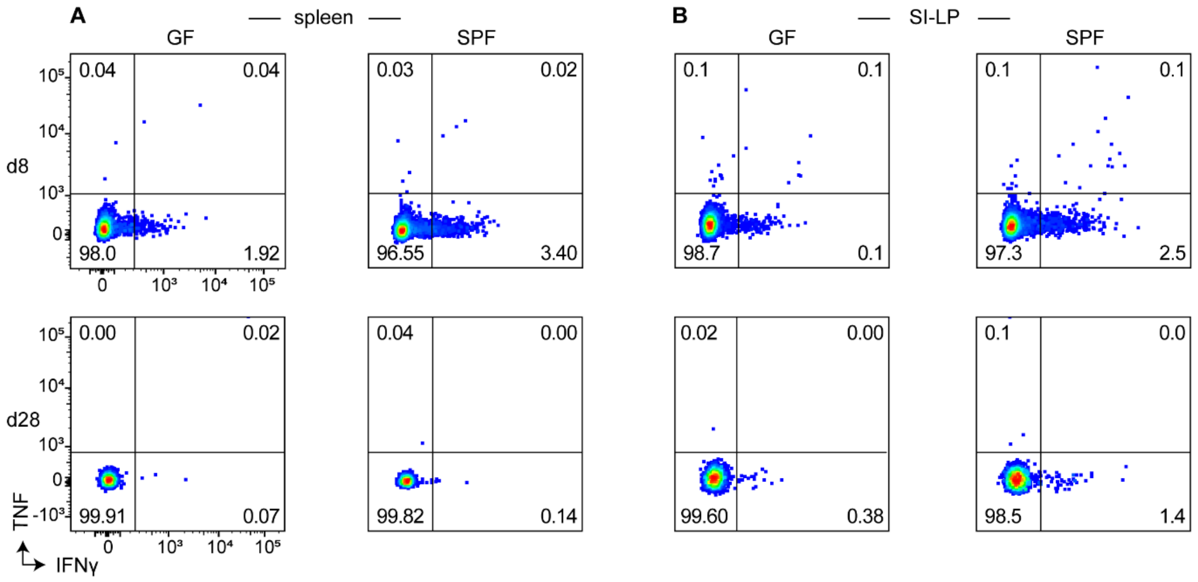


**Supp. Fig. 1: CD8 T cells in LCMV infected germ-free and SPF mice secrete cytokines specific to LCMV derived peptide after restimulation.** Experimental design: Germ-free (GF) and SPF C57BL/6 mice were infected with LCMV clone-13 and analyzed on day 8 or d28 after infection. **A** and **B**) Representative, CD8 gated flow cytometry plots showing the frequency of IFNγ and TNF expressing cells of GF and SPF mice on day 8 and d28 after infection in the spleen (A) and the SI-LP (B) after *in vitro* incubation in the absence of peptide.

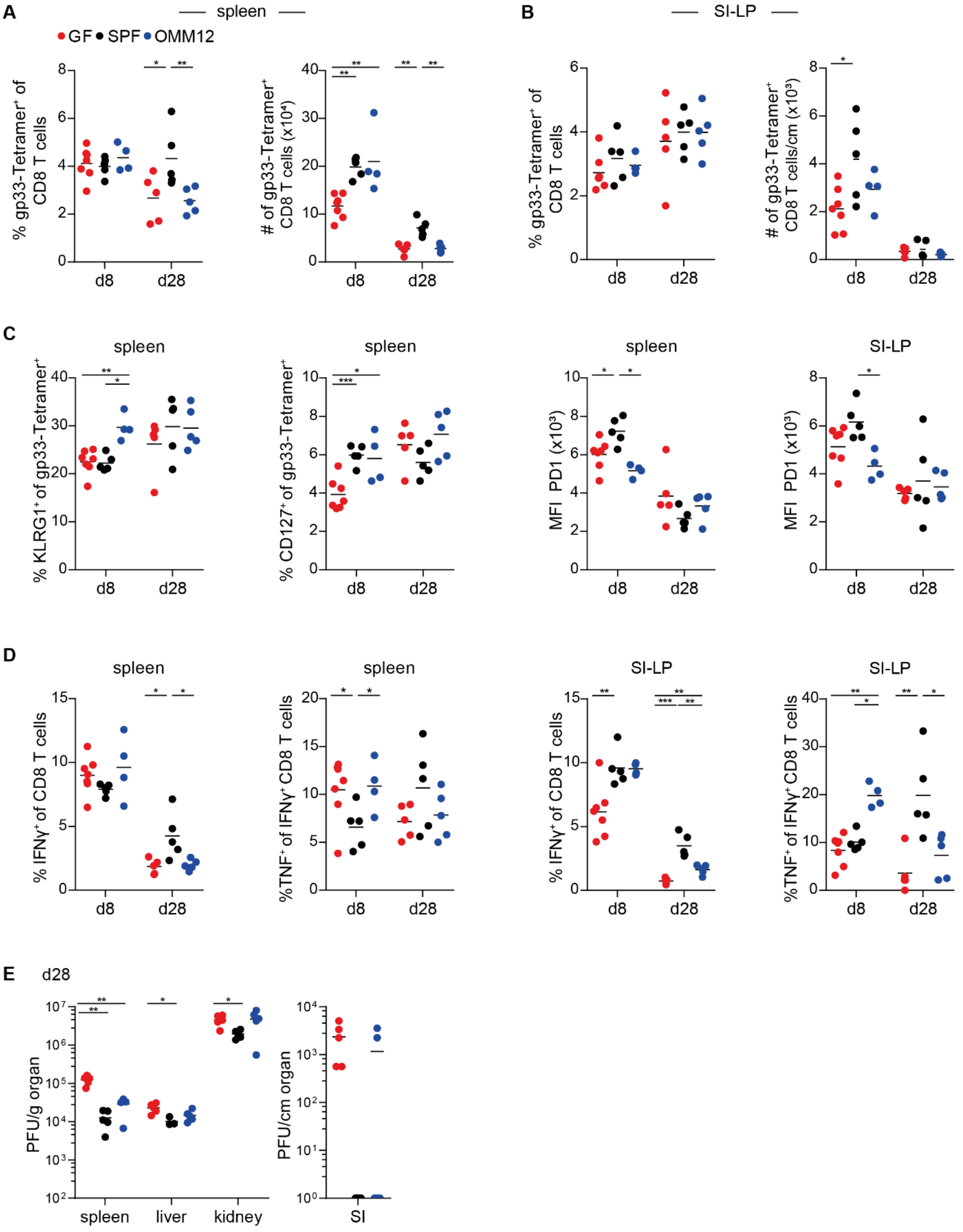


**Supp. Fig. 2: Similar phenotypic differentiation of LCMV-specific T cells in germ-free, SPF, and oligoMM-12 mice.** Experimental design: Germ-free (GF, red), SPF (black) and oligoMM-12 recolonized (OMM12, blue) C57BL/6 mice were infected with LCMV clone-13 and analyzed on day 8 or d28 after infection. **A** and **B**) Frequencies (A) and numbers (B) of tetramer+ cells of GF (red), SPF (black) and oligoMM-12 (blue) mice in the spleen and per centimeter of the SI-LP. **C**) Frequencies of KLRG1$^+$ and CD127$^+$ and MFI of PD-1 of tetramer$^+$ cells for GF, SPF and oligoMM-12 mice in the spleen (KLRG1, CD127 and PD-1) and the SI-LP (PD-1). **D**) Frequencies of IFNγ expressing (left) CD8 T cells and frequencies of TNF$^+$ of IFNγ expressing (right) CD8 T cells for GF, SPF, and oligoMM-12 mice. **E**) Viral titers in the spleen, liver, kidney (PFU/g), and SI (PFU/cm) were assessed on day 28 after infection. All data are from a single experiment with at least n ≥ 4; symbols indicate individual mice; horizontal lines show the mean; P values are from unpaired t-test or Mann-Whitney test (*p value < 0.05, **p value < 0.01, ***p value < 0.001).
